# Interdependence, Reflexivity, Fidelity, Impedance Matching, and the Evolution of Genetic Coding

**DOI:** 10.1101/139139

**Authors:** Charles W. Carter, Peter Wills

## Abstract

Genetic coding is generally thought to have required ribozymes whose functions were taken over by polypeptide aminoacyl-tRNA synthetases (aaRS). Two discoveries about aaRS and their tRNA substrates now furnish a unifying rationale for the opposite conclusion: that the key processes of the Central Dogma of molecular biology emerged simultaneously and naturally from simple origins in a peptide•RNA partnership, eliminating the epistemological need for a prior RNA world. First, the two aaRS classes likely arose from opposite strands of the same ancestral gene, implying a simple genetic alphabet. Inversion symmetries in aaRS structural biology arising from genetic complementarity would have stabilized the initial and subsequent differentiation of coding specificities and hence rapidly promoted diversity in the proteome. Second, amino acid physical chemistry maps onto tRNA identity elements, establishing reflexivity in protein aaRS. Bootstrapping of increasingly detailed coding is thus intrinsic to polypeptide aaRS, but impossible in an RNA world. These notions underline the following concepts that contradict gradual replacement of ribozymal aaRS by polypeptide aaRS: (i) any set of aaRS must be interdependent; (ii) reflexivity intrinsic to polypeptide aaRS production dynamics promotes bootstrapping; (iii) takeover of RNA-catalyzed aminoacylation by enzymes will necessarily degrade specificity; (iv) the Central Dogma’s emergence is most probable when replication and translation error rates remain comparable. These characteristics are necessary and sufficient for the essentially *de novo* emergence of a coupled gene-replicase-translatase system of genetic coding that would have continuously preserved the functional meaning of genetically encoded protein genes whose phylogenetic relationships match those observed today.

## Introduction: whence molecular genetics?

Gene expression consists of interpreting symbolic information stored in nucleic acid sequences. This irreversible computational process creates intrinsically novel meaning, and is thus fundamentally different from the physical chemistry underlying other natural processes, distinguishing it even from the molecular biological processes of replication and transcription. Our goal here is to provide a new conceptual basis for understanding how informational readout and the synthesis of peptide catalysts from instructions in genes might first have emerged and then evolved compatibly with inheritance.

### A. The Central Dogma and the adaptor hypothesis imply aminoacyl-tRNA synthetases (aaRS)

It is helpful to think about the origin of genetics within the conceptual framework first articulated by Crick with recent modifications (Fig. 1). Crick recognized that protein synthesis must be directed by information archived in DNA sequences and that information flow proceeds unidirectionally via an intermediate RNA “message” to ribosomes. He proposed separately that linking gene sequences to protein sequences required the intervention of a third RNA component (Crick 1955) to “adapt” individual amino acids to “codon” units in the message (Fig. 1A), accounting for the initially obscure relationship between what turned out to be collinear sequences of genes and proteins.

**Figure 1.**
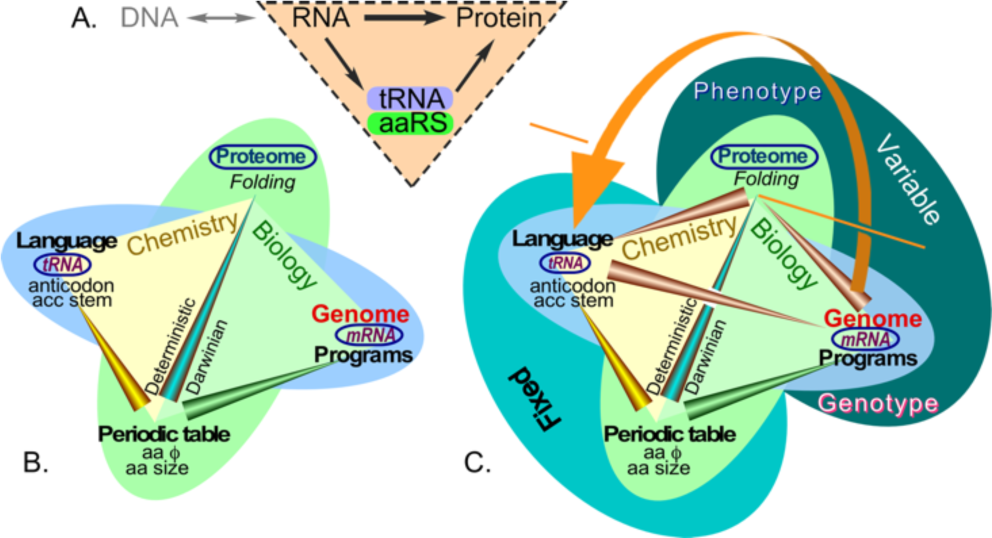
Information flow in molecular biology. A. The Central Dogma is supplemented by the “adaptor” hypothesis. The dashed triangle represents the crucial elements of Crick’s original insight, which necessarily implicates both tRNA and aaRS. B. The physico-chemical properties of the amino acids define the nano-scale “ecologies” within folded proteins, creating the intersection between genome and proteome. These ecologies also drove the selection of tRNA identity elements, analogous to a programming language, as well as protein folding. As a consequence, they also drive the selection of amino acid sequences in mRNA gene sequences (mRNA), analogous to computer programs. C. Network analysis of the Central Dogma consists of the nodes of a tetrahedron.Embedding the triangle from A into the ecology in B reveals a uni-directional feedback cycle or self-referential element as generator of complexity in the spirit of Gödel’s incompleteness theorem (Hofstadter 1979). Genetic instructions assemble amino acids according to their physical properties in ways that, when translated according to the programming language in tRNA, yield functional proteins (enzymes, switches, regulators). AARS with cognate tRNAs furnish reflexive elements (orange arrow) connecting their gene sequences, via their folded structures, to the enzymes that enforce the coding rules in the codon table. Physical properties of amino acids and the codon assignment table are “fixed” because they are governed by chemical equilibria. The genome and proteome are dynamically-determined biological processes that form the basis for the evolution of diversity through self-organization and natural selection of phenotypes.

Participation of the adaptor, transfer RNA (tRNA), involves creating a covalent bond between its 3’ terminus and the carboxylate group of an appropriate amino acid. Creation of that bond, in turn requires activation of the amino acid’s α-carboxyl group by reaction with ATP. In cells, activation and aminoacylation require a separate enzyme for each amino acid. These assignment catalysts, called aminoacyl-tRNA synthetases (aaRS), were first clearly identified by Berg and Ofengand (1958).

Executing genetic coding rules requires that aaRSs recognize both amino acids and tRNAs with high specificity so that the former can be escorted to the ribosome by the latter for protein synthesis. However, specific recognition by folded proteins depends on a complex “ecology” based on the chemical behavior and interactions of individual amino acids (Fig. 1B). That behavior can be accurately parameterized by two experimental Gibbs phase transfer free energies—from vapor to cyclohexane and from water to cyclohexane—related to the size and polarity, respectively, of each amino acid’s side chain (Carter and Wolfenden 2015; Wolfenden, et al. 2015; Carter and Wolfenden 2016). Correlations between these free energies and tRNA identity elements recognized by aaRS and the distribution of amino acids between surfaces and cores after protein folding established these parameters as the main axes of a kind of “periodic table” of amino acids (Carter and Wolfenden 2016) concatenated in chains that fold to generate proteins of virtually unlimited functional diversity, in analogy to joining atoms to form molecules.

Implementing the Central Dogma—the irreversible attachments of amino acids to codon-specific tRNAs by aaRSs—thus exploits the ecology of the amino acids within those enzymes. Proteins folded in accordance with such ecologies that, in turn, execute computationally controlled production from genes of specialized amino acid ecologies (including their own!) compose a reflexive property known as a “strange loop” (Fig. 1C; (Hofstadter 1979). Recognizing that loop opens fundamentally new ways to think about what enabled the aaRS to emerge as the only proteins coded by programs written as mRNA that can, once folded, collectively interpret the programming language in tRNA. We propose that this reflexivity in functional chemistry and encoded information played a crucial role in creating genetics.

### B. The RNA World hypothesis fails to address key questions about gene expression

The default framework for thinking about how genetics emerged has been a facile solution to the problem that life simultaneously requires that genetic information must be passed from generation to generation, and that catalysts must synchronize rates of chemical reactions underlying the accuracy in gene replication, expression, and metabolism. Base pairing between complementary nucleic acid strands answered the former problem immediately and decisively, once the helical structure of double-stranded DNA was elucidated (Watson and Crick 1953), and pointedly highlighted the second problem.

The crystal structure of tRNA^Phe^ (Kim, et al. 1973) revealed that, unlike DNA, RNA can assume tertiary structures, consistent with proposals (Woese 1967; Crick 1968; Orgel 1968) that the earliest catalysts also might have been RNAs that could “do the job of a protein” (Crick 1968). That hypothesis has been sustained almost exclusively by the observation that, whereas proteins cannot readily store or transmit digital information, RNA does have rudimentary catalytic properties (Cech 1986; Guerrier-Takada 1989). The expedient conclusion that RNAs functioned as both genes and catalysts in a life form devoid of proteins was rapidly embraced as “the RNA World” (Gilbert 1986).

The clarity with which base-pairing solved the inheritance problem and the discovery of, and fascination with, catalytic RNA short-circuited the quest to understand and answer deeper questions:

*Catalytic RNA itself falls far short of fulfilling the tasks now carried out by proteins.* The term “catalytic RNA” overlooks three fundamental problems: (i) it vastly overestimates the potential catalytic proficiency of ribozymes (Wills 2016); and fails to address either (ii) the computational essence of translation, or (iii) the requirement that, throughout the evolution of translation and intermediary metabolism, catalysts not only accelerate, but more importantly, synchronize chemical reactions whose spontaneous rates differ by than 10^20^–fold (Wolfenden and Snider 2001).

*The nexus connecting pre-biotic chemistry to biology is not replication but the translation table that maps amino acid sequences of functional proteins onto nucleotide triplet codons*. The quintessential problem posed by life’s diversity (Carter and Wolfenden 2016; Wills 2016) is how that critical transformation became embedded, in parallel, into tRNA and gene sequences, together with the ribosomal read-write mechanism (Bowman, et al. 2015; Petrov and Williams 2015). Spontaneous folding of RNA aptamers and the dynamics of an RNA world do not require encoding into genetic information and hence fall well short of what “conversion to a functional molecule” (Horning and Joyce 2016) implies for Darwinian evolution by selection acting on phenotypes (Wills 2016).

*By the time protein folding organizes amino acid side chains into a functional active site, genetic information has been irreversibly transformed.* Any molecular machine charged with reversing translation by unfolding, then “reading” the sequence of a protein would require shuttling each successive amino acid through ~20 active sites until one fitted, and then overcoming the redundancy of the genetic code. By enabling the inheritance of genetically encoded characteristics this one-way flow of genetic information enshrined in the Central Dogma (Koonin 2015) ensures that biological evolution transcends the simple population dynamics of natural selection in any RNA world.

*RNA research has never provided, either experimentally or conceptually, even an approximate model for how a nearly random catalytic network, without encoded proteins, might have progressively bootstrapped the specificity and selectivity characteristic of enzymic systems (Hordijk, et al. 2014).* Thus, evolution of synchronized catalysis required simultaneous evolution of genetic coding.

### C. Many nevertheless embrace the RNA World with little reservation (Wolf and Koonin 2007; Van Noorden 2009; Yarus 2011b, a; Bernhardt 2012; Breaker 2012; Robertson and Joyce 2012)

It is important, therefore, to assess the experimental data on which the hypothesis rests and to separate data that genuinely support the hypothesis from those that only appear to do so.

*Selecting ever more proficient RNA aptamers from large combinatorial libraries based originally on self-splicing introns only appears to support the existence of ancestral ribozymal polymerases*. Despite the technical elegance and practical value of Selex experiments (Tuerck and Gold 1990), even fully-developed aptamer replicases (Wochner, et al. 2011; Attwater, et al. 2013; Sczepanski and Joyce 2014; Taylor, et al. 2015; Horning and Joyce 2016) would support only a limited version of the RNA World hypothesis without phylogenetic relationships connecting them to biological ancestry. So far as we know, all nucleic acids in contemporary biology are synthesized by protein enzymes, much as, reciprocally, the synthesis of proteins from activated amino acids is catalyzed by an RNA template at the peptidyl transferase center of the ribosome (Noller, et al. 1992; Noller 2004; Petrov, et al. 2014; Bowman, et al. 2015). Thus no phylogenetic basis exists for ancestral ribozymal polymerases.

*The most compelling evidence that proteins were first coded by ribozymes is the extensive phylogenetic analysis of contemporary protein families.* Koonin and colleagues (Aravind, et al. 2002; Leipe, et al. 2002; Koonin and Novozhilov 2009; Koonin 2011), and others (Caetano-Anolles, et al. 2007; Caetano-Anollés, et al. 2013; Caetano-Anollés and Caetano-Anollés 2016) argue that protein domains speciated substantially before the advent of protein-based aminoacyl-tRNA synthetases and translation factors. Consequently, they argue, a fully developed ribozyme-based version of the contemporary universal genetic code must have first mapped RNA sequences to the amino acid sequences of peptides. We will call this fully-blown RNA World scenario the “RNA Coding World” (RCW; see also (Rodin and Rodin 2006b, a; Rodin and Rodin 2008; Rodin, et al. 2011)). The contemporary “Protein Coding World” (PCW), which uses aaRS enzymes to attach amino acids to cognate tRNAs, is envisaged to have evolved by a series of “takeovers”, whereby the coding functions of aaRS ribozymes were progressively replaced, without disruption, by enzymic counterparts. Our analysis articulates the improbability that such a takeover could ever have taken place.

### D. Contemporary aaRSs furnish clues about how they became molecular interpreters

Understanding the evolutionary basis for the Central Dogma (Fig. 1) requires asking how self-organization and selection might have produced, from nearly random origins, finely tuned ecological niches of amino acids arranged to provide the catalytic and pattern-matching capabilities necessary for the operation of a code using a 20-letter alphabet. We envision that this process began with a reduced alphabet administered by a small “boot block” that grew by stepwise increases in alphabet size, in which the information that survived (i.e. was selected) at each stage was such that it could be used by the existing interpreters to make themselves, in spite of the errors that they made.

Accumulating such errors leads potentially to two types of “catastrophes”. Replicative errors eventually limit the survival of progressively longer “genes”, and can produce what has been called an “Eigen catastrophe” (Eigen and Schuster 1977). Similarly, translation errors eventually limit the functional specificity available to maintain a cell’s biochemical network, and can lead to an “Orgel catastrophe” (Orgel 1968). Thus, replication and translation errors represent the most significant resistance to the emergence and gradual enhancement of biological complexity.

Eigen (Eigen 1971; Eigen and Schuster 1977) articulated a strategy for integrated survival of both information and functional specificity in an error-prone network involving “information carriers” and “functional catalysts”. He noted that the survival of separate molecular species may be enhanced in systems containing either codependent or multiply interdependent components, whose cooperation with other members of the set might assist survival of sets of molecules that would otherwise be eliminated by competition. He called the symmetrical arrangement of components within these sets “hypercycles”, a concept that can be generalized to include other interdependent arrangements.

Although early aaRS phylogenies should record the order in which enzymic aaRS appeared, either *ab initio* or during their takeover of ribozymal aaRSs, earlier authors (Woese, et al. 2000) doubted that such information would be relevant to the emergence of genetic coding. §I, however, summarizes a new interpretation of evidence, from experimental deconstruction of both aaRS classes (Chandrasekaran, et al. 2013; Carter 2014; Carter, et al. 2014; Carter 2015, 2016, 2017), that all contemporary aaRS descended in modular fashion from a single bi-directional gene, whose strands coded for functional ancestors, respectively, of Class I and II synthetases. Products of that gene appear to have been optimally differentiated and crafted almost ideally to establish hypercycle-like interdependence, implementing a minimal amino acid alphabet—all characteristics of the “boot block” envisioned to have first enabled genetic coding. This bi-directional coding ancestry necessarily coupled the evolutionary descent of contemporary Class I and II aaRSs phylogenies (O’Donoghue and Luthey-Schulten 2003; Wolf and Koonin 2007; Caetano-Anollés, et al. 2013) discussed in §B above.

We therefore face a stark choice: either the sequential aaRS decompositions into increasingly conserved fundamental modules—urzymes (Ur = primitive; (Li, et al. 2013; Carter 2014; Martinez, et al. 2015)) and protozymes (Proto = before; (Martinez, et al. 2015)—and their bi-directional coding ancestry (Chandrasekaran, et al. 2013; Carter, et al. 2014; Carter 2015), or the relative origins of multiple protein superfamilies derived from current phylogenetic analyses must be wrong. We outline a resolution in §III. Phylogenetic and biochemical evidence has been supplemented by developing statistically significant (Carter and Wolfenden 2015, 2016) relationships between identity elements that dictate recognition of tRNA by different synthetases and the size and polarity of amino acid sidechains. Coding relationships implemented in tRNA recognition are therefore not arbitrary, but reflect the deeply relevant inner logic of protein folding rules (Carter and Wolfenden 2015; Wolfenden, et al. 2015; Carter and Wolfenden 2016). We consider this point, reflexivity, and other relevant concepts in greater detail in §II.

Correlations of the tRNA identity elements also revealed that signals for the two physical properties are distributed differently between the anticodon (specifying polarity) and the acceptor stem (specifying size). This, in turn, furnished details of an “operational RNA code” in the tRNA acceptor stem (Schimmel, et al. 1993) that likely preceded development of coding by the tRNA anticodon, perhaps first implementing only the binary discrimination between large (Class I) and small (Class II) side chains. This new evidence makes it difficult to imagine how a sophisticated proteome synthesized by an advanced ribozyme-based translation system could have survived any transition via a protein-based expression system with fidelity as low as that of primordial binary coding.

## RESULTS

### I. AARS class dualities would have helped to stabilize quasispecies bifurcations

Three functionalities give the aaRS their unique status as the earliest enzymes: (i) They accelerate amino acid activation at the expense of two ATP phosphates ~10^14^-fold, resulting in irreversible synthesis of aminoacyl 5’AMP. The uncatalyzed rates of all other reactions in protein synthesis are orders of magnitude faster than that reaction, which thus limits the rate of prebiotic protein synthesis. (ii) The adenosine in ATP serves as an affinity tag that increases active-site binding 1000-fold, enhancing coding assignment specificity, especially where editing is required. (iii) They acylate tRNA, covalently linking a specified amino acid to a tRNA molecule bearing a code-cognate anticodon.

Notably, two distinct sets of homologous aaRS structures, Class I and Class II (Cusack, et al. 1990; Eriani, et al. 1990; Ruff, et al. 1991), implement these three functions in disparate ways. The two classes activate symmetrical sets of 10 amino acids. Both classes have one major (A), and two different minor subclasses (B and C) (Cusack 1994). The common origin of the two aaRS classes on opposite strands of the same ancestral gene (Rodin and Ohno 1995) remained obscure until quite recently (Martinez, et al. 2015). This section outlines how consequences of this duality at multiple structural and functional levels may have served to differentiate and stabilize early stages of genetic coding in the face of high error rates.

*An ancestral bi-directional gene produced a Class I ancestor with a modest specificity for larger amino acids, and a Class II ancestor with similar specificity for smaller amino acids.* No useful information can be encoded using only one kind or equivalent class of activated amino acid. The simplest imaginable code would have required discriminating between at least two kinds of amino acids. The interesting scenarios (Wills 2004) thus entail generating the full code from simple 2- or 4-letter alphabets via transitions, each of which increases the effective size *n_eff_* of the amino acid and codon alphabets. Nested instabilities (Wills 2004) allow for such a series of code-expanding transitions to attractor states with progressively larger values of *n_eff_*. These transitions connect dynamic states with significant error rates and thus entail broad distributions of functional protein sequences and their encoding genes that are called “quasispecies”, so we call the corresponding transitions “quasispecies bifurcations” (Fig. 2).

**Figure 2.**
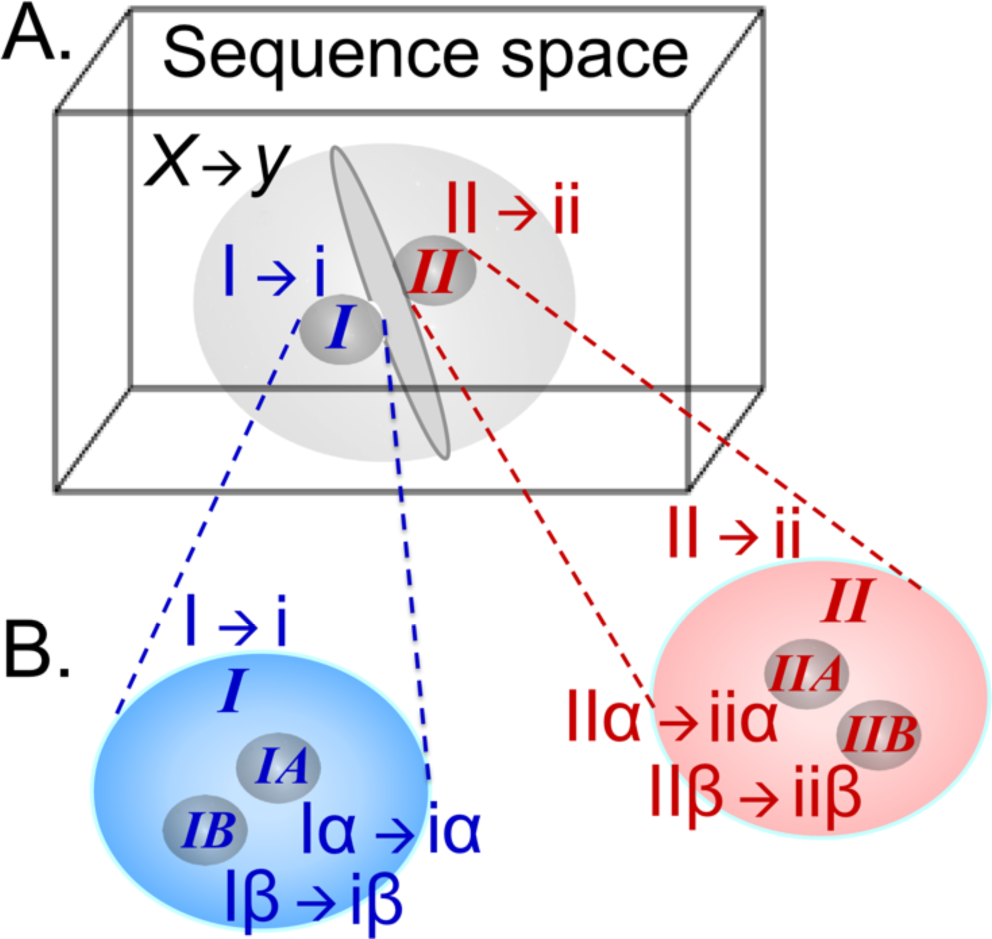
Quasispecies bifurcations in aaRS gene or protein sequence space. A. A single quasispecies of undifferentiated function making random assignments *X* → *y* of codons (*X*) to amino acids (*y*) cannot transmit genetic information. Nor can it easily bifurcate to a pair of narrower quasispecies. Bi-directional coding ancestry of the contemporary aaRS created suitable quasispecies *de novo* {***I*, *II***; red and blue; bold italics explicitly indicating Class I, II aaRS} each separately supporting binary coding assignments I → i and II → i of specific subsets of codons {I, II} to corresponding subsets of amino acids {i, ii}. That double-helical gene with dual single-strand interpretations overcame the initial and most substantial barrier to the emergence of genetic coding by partitioning protein sequence space decisively into two functionally distinct populations. The plane between the ***I*** and ***II*** quasispecies is a local representation of the inversion operator that transforms a sequence into its complement read in the reverse direction. B. Daughter population distributions derived from nearly simultaneous bifurcation of the two ancestral binary coding quasispecies (expanded view) into smaller separate sub-populations of genes and assignment catalysts operating a 4-letter code {Iα → iα, Iβ → iβ, IIα → iiα, IIb → iiβ,}. Genetic coding bi-directionality is preserved through complementary gene pairs Iα⇔IIα and Iβ⇔IIβ. Recapitulation of the bifurcation process would further specialize related species, each step being progressively easier owing to the increased coding specificity, but eventually losing the ability to use information in both strands of genes.

The TrpRS and HisRS urzymes (Li, et al. 2013) and the designed Class I/II protozyme gene (Martinez, et al. 2015) furnish substantive experimental representations of the ancestral assignment catalysts envisioned by Wills (2004). Gene products created from opposite strands utilizing the full genetic code both accelerate amino acid activation ~10^6^-fold. Both wild-type protozymes exhibit high ATP affinity and the Class I protozyme possesses a consensus phosphate binding site composed entirely of oriented backbone NH groups (Hol, et al. 1978). Thus, it seems plausible and of obvious interest that protozymes coded using fewer than the canonical 20 amino acids might retain substantial catalytic activity.

A coding system assigning dual classes of amino acids {α,β} that are functionally differentiated in a crude binary fashion to tRNAs with anticodons complementary to codons {A,B} by such species could bifurcate into two versions to produce four-member codon and amino acid alphabets, {A, B, X, Δ} and {α, β, χ, δ}, increasing the coding capacity from 2 letters to 4 letters, and expanding the 2 × 2 translation table into a 4 × 4 table. In simulations of such a process (Wills 2004, 2009), the hierarchically nested embedding of assignment activities in the protein sequence space geometrically mirrored the decomposition of the alphabets. The system showed stepwise coding self-organization, first from a non-coding state to the execution of a binary code {A→a, B→b} and then from the binary code to the expanded four-dimensional code {A→α, B→β, X→χ, Δ→δ}, anticipating experimental studies of the two synthetase Classes (Fig. 2). Ancestral bi-directional coding would have impacted the emergence and evolution of genetic coding in several important ways. First, the ancestral gene would have partitioned the sequence space decisively, dividing it between those sequences related most closely to each of the two strands. Second, the translated products of each strand would have differentiated the functional specificities retained by sequences surrounding the centroids of the two populations. Third, the bidirectional coding complementarity constraint steepens the fitness landscape, decisively enforcing coding cooperation compared to the corresponding possibilities for genes that could mutate independently, thereby increasing selection pressure for coding. Cooperation is therefore more robust than it would be with independent genes for the Class I and II urzymes. Finally, the reduced volumes of sequence space and enhanced functional specialization of the two bi-directionally coded quasispecies suggest that fewer mutations were necessary for neofunctionalization of subsequent duplications, successively easing subsequent bifurcations as *n_eff_* increased during the bi-directional coding regime.

#### A. Experimental deconstructions of Class I and II aaRS reveal parallel structural hierarchies

A puzzling hierarchy of inversion symmetries in the structural, functional, and evolutionary biology of contemporary aaRSs now appears to make sense if the aaRSs are remnants of such bifurcations.

*Superimposing Class I and II aaRS catalytic domains reveals small invariant cores, distinct from idiosyncratic elements unique to each amino acid.* Like Russian Matryoshka dolls, parallel deconstruction of both Class I and II aaRS families reveals nested, increasingly conserved modular catalysts of nearly equal molecular mass (Carter 2014): catalytic domains (200-350 residues), urzymes (120-130 residues; (Pham, et al. 2007; Pham, et al. 2010; Li, et al. 2011; Li, et al. 2013)), and protozymes (46 residues; (Martinez, et al. 2015)), each retaining conserved portions from its preceding construct.

Urzymes retain all necessary functions of full-length aaRS, albeit with lower proficiency and specificity, and are analogous to using “molecule” to define the smallest unit of matter that retains all properties of a chemical substance. Protozymes, on the other hand, approach the smallest polypeptide catalysts and hence are perhaps more analogous to “atoms”.

Published evidence that experimental urzyme catalytic activities arise neither from tiny amounts of wild-type enzyme nor from unrelated, but highly active contaminants includes the following (Pham, et al. 2010; Li, et al. 2011): (i) empty vector controls have no activity. (ii) protease cleavage tagged fusion proteins releases cryptic activity. (iii) Mutations alter activity. (iv) Amino acid *K_M_* values differ from WT values, and, most importantly, (v) single turnover active-site titration experiments show pre-steady-state burst sizes demonstrating that 35−75% of molecules transiently form tight transition-state complexes. Experimental assays of protozymes were validated by showing that active-site mutants H18A (Class I) and R113A (Class II) eliminated activity of the respective catalyst (Martinez, et al. 2015).

*Modular accretions in the structurally unrelated Class I and II protein superfamilies exhibit parallel accelerations of the rate-limiting step of protein synthesis over a 10^8^-fold range.* Experimental transition-state stabilization free energies track linearly with number of residues in deconstructed constructs from both Classes, reinforcing the justification of these constructs as snapshots in the parallel evolution of both synthetase classes (Martinez, et al. 2015). Urzymes retain ~60% of the full-length transition state stabilization free energy observed in modern synthetases. Protozymes are only 46 amino-acids-long. Although they retain only the ATP binding sites, aaRS protozymes from both Class I and II aaRS exhibit ~40% of the full-length transition-state stabilization, but have not yet been shown either to acylate tRNA or to discriminate significantly between different amino acids.

These accelerations document that multiple families of proteins are capable of synchronizing chemical reactions over a very broad range from the uncatalyzed rate to that observed in contemporary organisms. RNA has not been shown capable of parallel rate accelerations over such a dynamic range either in parallel families or with similar increases in mass, underscoring the superior ability of polypeptide catalysts to synchronize cellular chemistry.

#### B. Ancestral bi-directional genetic coding underlies the aaRS class distinction

Rodin and Ohno (Rodin and Ohno 1995) aligned coding sequences of the two aaRS Classes in opposite directions revealed highly significant bi-directional coding of the class-defining active-site sequence motifs. Subsequently, it became increasingly apparent that protein-based aaRSs all descended from a single ancestral gene whose complementary strands encoded precursors to the Class I and Class II aaRS superfamilies (Carter, et al. 2014; Carter 2015; Martinez, et al. 2015). Bi-directional coding ancestry implies that protein aaRS gene evolution began with an early stage in which the unique information in one strand of a gene could be interpreted as a different protein with a similar function on the opposite strand. Three types of results have confirmed predictions of the Rodin-Ohno hypothesis:

1) The most highly conserved portions of contemporary aaRSs should correspond to modules in the contemporary enzymes capable of bi-directional alignment, and should retain catalytic activity when extracted from the full-length genes. Two successive levels of experimental deconstruction confirm this prediction. Urzymes (Pham, et al. 2007; Pham, et al. 2010; Li, et al. 2011; Li, et al. 2013) have ~120-130 amino acids and retain all of the translation functions of contemporary synthetases and accelerate amino acid activation by 10^9^-fold, with significant specificity. A designed bi-directional gene encodes ~46 amino acid Class I and II Protozymes that contain the ATP binding sites of the respective aaRS, bind ATP tightly, and accelerate amino acid activation 10^6^-fold (Martinez, et al. 2015).

2) Coding sequences should retain a higher frequency of base-pairing between middle codon bases in antiparallel, in-frame alignments of Class I and II aaRS. This middle-base pairing frequency, ~0.34, is significantly non-random and increases to 0.42 in comparisons between coding sequences reconstructed independently for ancestral nodes of both Class I and II aaRS (Chandrasekaran, et al. 2013).

3) It should be possible to re-construct a *bona fide* bi-directional gene such that each strand codes for a functional amino acid activating enzyme homologous to one contemporary aaRSs class. We configured Rosetta to both constrain tertiary structures and impose genetic complementarity to give “designed” Class I and II protozymes (Martinez, et al. 2015). Remarkably, all four wild-type and designed peptides from Class I and Class II have the same *k_cat_/K_M_* and accelerate amino acid activation by ~10^6^-fold. Wild-type sequences have 100-fold lower *k_cat_* and 100 fold higher *K_M_* values than do the designed protozymes from the complementary gene, in keeping with the possibility that their wild-type sequences may include amino acid binding determinants lost in the designed protozymes. The protozymes extend a linear relationship between transition state stabilization free energy and the number of residues of the constructs. Notably, Class I and II constructs exhibit the same slopes and intercepts relating rate acceleration to number of residues (Martinez, et al. 2015).

Bi-directional, in-frame coding is a strange idea. Base-pairing is part of an inversion symmetry operator that generates the *sequence* and (using helical symmetry operators) the *structure* of the opposite strand. Because the opposite strand sequence can be retrieved using this inversion operator, a gene’s unique information is contained in each strand. That unique information, however, has two different functional interpretations. Validating (1)-(3) of the Rodin-Ohno hypothesis revealed higher-order symmetries relating Class I and II gene products (Carter, et al. 2014; Carter 2015), as discussed in §II.C - §II.E.

#### C. An ancient hypercycle-like interdependence relates catalytic residues in each Class

Active-site amino acids in aaRS occur in three sets of signature sequences (Eriani, et al. 1990; Carter 1993). Class I HIGH and KMSKS sequences and Class II Motifs 1 and 2 are present in the respective urzymes. The HIGH/Motif 2 signature is present in the protozymes. As these motifs provided the original evidence for bi-directional coding (Rodin and Ohno 1995), and contain active-site residues, it comes as no surprise that the respective active-sites utilize different catalytic residues. In fact, all residues contributing to catalysis by Class I active sites must be activated by Class aaRS II, and conversely, residues needed for Class II activity must be activated by Class I aaRS (Carter, et al. 2014; Carter 2015, 2017). This functional “anti-homology” dates from the earliest Class I and II catalysts. Interdependence induces a hypercycle-like coupling between the two bi-directional gene products similar to that proposed by Eigen (Eigen 1971; Eigen and Schuster 1977) to mitigate competition, induce cooperation and thereby increase the overall semi-random genetic content that could survive deterioration at any given copy-error rate.

#### D. Folded Class I and II AARS tertiary structures are “inside out”

Binary patterns coding for protein secondary structures (Kamtekar, et al. 1993; Patel, et al. 2009) are reflected across complementary coding strands. They are determined by positions of hydrophobic residues (Muñoz and Serrano 1994); the heptapeptide repeat, a-g, with hydrophobic amino acids in positions a,e,f, is diagnostic for alpha helix. Alternation of hydrophobic side chains, especially when they include side chains with branched β-carbon atoms, is almost as effective a predictor of β-structure.

Soluble globular proteins have hydrophobic cores and water-soluble surfaces. The distribution of amino acids in folded proteins between these two extreme environments is spanned by a two-dimensional “basis set” furnished by the experimental free energies of transfer between vapor and cyclohexane and between water and cyclohexane (Carter and Wolfenden 2015; Wolfenden, et al. 2015). The contemporary genetic code respects this dichotomy to an extraordinary degree, as codons for virtually all core side chains are anticodons for distinctly surface side chains (Zull and Smith 1990). Complementary codons for proline and glycine, most often associated with turns, mean that such sequence-directed turn formation also reflects across codes from antiparallel strands. Thus, the folded products from a bi-directional gene will tend to have comparable secondary structures, with opposite polarities. By these criteria, Class I and II aaRS urzymes are both antiparallel and “inside out”.

#### E. Class I and II aaRS amino acid substrate specificities, especially those from ancestral codes, are related by inversion with respect to side chain size (Carter, et al. 2014; Carter 2015)

Modern aaRSs prefer their cognate amino acids by ~5.5 kcal/mole, ~80% of which comes from allosteric influences of more recently acquired modules on the urzyme activities. Lacking insertion- and anticodon-binding domains, Class I LeuRS and Class II HisRS urzymes are relatively non-specific (Carter, et al. 2014; Carter 2015). Experimental Δ*G_kcat/KM_* values show that they have *similar and complementary specificities*. LeuRS urzyme prefers Class I substrates; HisRS urzyme prefers Class II substrates, both by ~ 1 kcal/mole. They are therefore capable of making the correct choice between Class I and II amino acids roughly four times in five. That fidelity is too promiscuous to support more than “statistical ensembles” of peptides, as hypothesized by Carl Woese (Woese 1965a, b; Woese, et al. 1966). Thus, urzymes would have been the predominant assignment catalysts within a much broader population of molecular types, with the properties of a “quasispecies-like” cloud as defined by Eigen (Eigen and Schuster 1977) that likely included many species with similar specificity, but lower catalytic efficiency.

The only statistically significant distinction between amino acids activated by Class I and Class II aaRS is their sizes (Carter and Wolfenden 2015; Wolfenden, et al. 2015): Class I amino acids are uniformly larger than those from Class II. Accounting for the solvent exposure of amino acids in folded proteins entails both size and polarity and is therefore two-dimensional (Carter and Wolfenden 2015, 2016). Class II amino acids migrate significantly toward water interfaces during protein folding, whereas Class I amino acids migrate toward cores. This differential localization in folded proteins gains significance in the context of the distribution of identity elements in tRNA (§II.F).

#### F. tRNA acceptor stem identity elements represent a code for amino acid side-chain size and other descriptors including side chain carboxylation and β-branching

Evidence that the much smaller aaRS urzymic cores could have accelerated the rates of tRNA aminoacylation (Li, et al. 2013) now makes it increasingly likely that an early “operational genetic code” (Schimmel, et al. 1993; Schimmel 1996) functioned entirely on the basis of acceptor stem bases that specified the most significant difference between Class I and Class II amino acids. Ancestral tRNAs may have been only about half the size and consisted of only the acceptor and TΨC loops of modern tRNAs. Doubling of this ancestral structure has been proposed to have created the anticodon and dihydrouridine loops with the anticodon initially serving as a proxy for the identity elements in the acceptor stem (Di Giulio 1992; Rodin, et al. 1996; Di Giulio 2004, 2008; Rodin and Rodin 2008). Any successful model for the emergence of genetic coding from an RNA-based system of molecular information processing should thus be consistent with these two observations as well as with the phylogenies of the two aaRS Classes.

#### G. Class I, II genes, gene products, mechanisms, and specificities are maximally differentiated

An important barrier to the emergence of diversity from quasi-random reproductive processes is the strong tendency of mutant daughter species to regress to the centroid of the distributions from which they originate (Eigen, et al. 1988). The centroids behave as “strong attractors”. Inversion symmetries relating Class I and II aaRS, described in §II.B-§IIE suggest that their genes, gene products, functions, and substrates are inherently differentiated to survive successive quasispecies bifurcations necessary for enhanced genetic coding to emerge from populations of low sequence identity and modest specificity:

1. Bi-directional coding complementarity means that individual ancestral Class I and II gene sequences are as difficult as possible to interconvert from one to the other by serial mutation.
2. Descent of the Class I and II aaRS from a bi-directional gene stabilizes two quasispecies that can presumably begin to interpret binary sequence patterns, decisively overcoming the barrier posed by the strong attraction of a single quasispecies.
3. Reduced population size and enhanced functional specialization of the two bi-directionally coded quasispecies suggest that fewer mutations are necessary for neofunctionalization, successively easing subsequent bifurcations during the bi-directional coding regime.
4. Distinct properties of protozymes and urzymes point to successive emergence during the bi-directional coding era of their ATP-, amino-acid- and pyrophosphate-binding sites, consistent with modular construction of basic aaRS functions.
5. Inverted folding instructions give rise to “inside out” Class I and II tertiary structures that are as different as possible from one another, and thus minimally vulnerable to mutations that might fuse the two quasispecies by regression to the common centroid.
6. Catalytic residues in Class I and II aaRS are entirely segregated. Thus, throughout their early evolution, the two Classes formed a hypercycle-like network (Fig. 3). By arguments from Eigen and Schuster (Eigen, et al. 1988) and Wills (Wills 2009), their interdependence defended them against corruption by molecular parasites during growth of catalytic networks.
7. Class I and II amino acids are themselves optimally separated on the basis of (i) size, (ii) polarity, and hence (iii) their ultimate destination in folded proteins.

**Figure 3.**
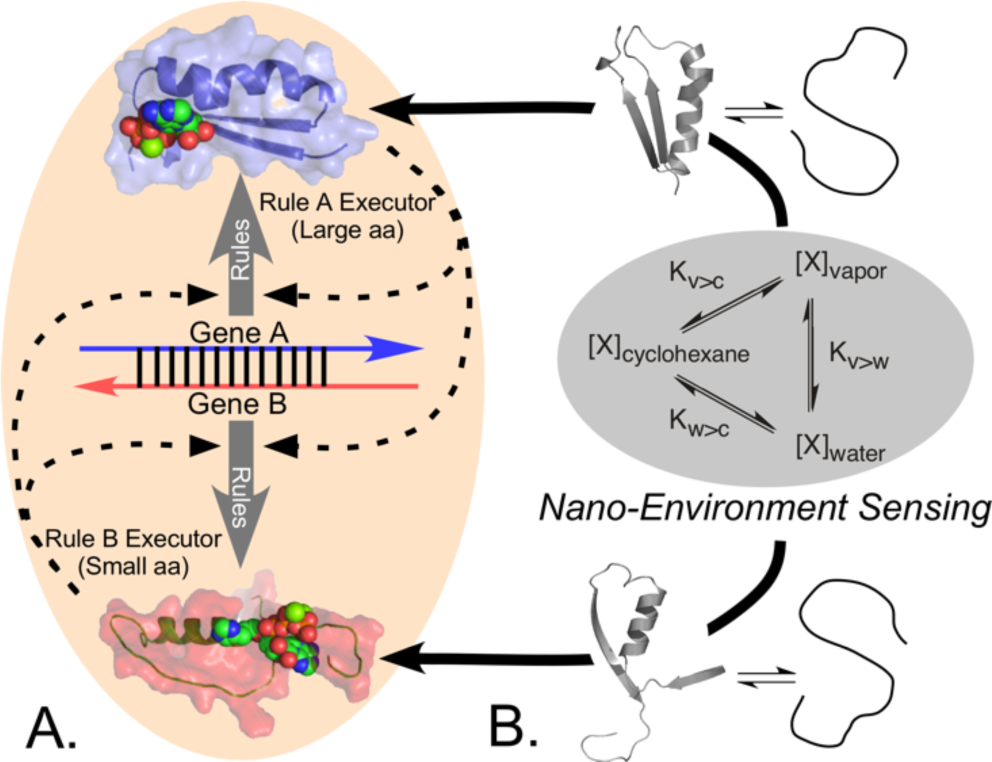
Reflexivity is an exclusive property of protein aaRS that arises from nano-environmental sensing. A. The putative ancestral amino acid activating protozyme, substantiated experimentally in (Martinez, et al. 2015) furnishes two assignment catalysts, each executing a complementary assignment, one for large, the other for small amino acid sidechains. Each also contributes to the translation of the other. B. As the assignment catalysts in A are proteins, their folding reactions are governed by the phase transfer equilibria of the amino acids, sensing the nano-environment in a necessary prelude to function.

### II. Bi-directionality furnishes four properties indispensable for self-organization of coding

Avoiding multiple stop codons on both strands of a bi-directionally coded ancestral gene would mandate that each of the four bases have a functionally coded meaning when it occurs as an (internal) codon middle base (see for example (Delarue 2007)). This would imply a (possibly redundant) alphabet of four letters. Such a reduced repertoire is consistent with that expected for an ancestral tRNA acceptor stem, in keeping with the fact that the contemporary acceptor stem code distinguishes best between (i) large and small, (ii) β-branched vs unbranched, and (iii) carboxylate vs non-carboxylate side chains (Carter and Wolfenden 2015). Presumably, selection subsequently drove both the code and primordial coding sequences to capture and employ additional symbolic information for precisely those chemical properties—size and polarity—that determine how the 20 amino acids direct proteins into unique configurations (Carter and Wolfenden 2016).

Bi-directional coding of enzymic aaRS impacts four properties that favor much more rapid and efficient evolution of gene expression than would have been possible for ribozymal aaRS. These properties are developed more extensively and with greater mathematical rigor in a separate paper (Wills and Carter 2017).

#### A. Any set of aaRSs forms an interdependent catalytic network

Because contemporary aaRS are proteins, their own functional structures all depend intimately on all aaRS functionalities and so form hypercycle-like networks. Interdependence implies further that both programming language in tRNA and their mRNA programs co-evolved from simpler ancestors with fewer distinctions between them, and hence less complex interdependencies. Therefore, they are expected to have something approaching discrete ancestries, and hence successively simpler levels of interdependence as we approach the root. Structural variants in any functional aaRS population must have responded coordinately throughout their evolutionary history to two different chemical signals—amino acid and tRNA. Bi-directional coding ancestry deeply anchors such interdependence in the earliest ancestral quasispecies, as active-site catalytic residues in Class I and II aaRS must be activated by the opposite class (Carter, et al. 2014; Carter 2015, 2017).

#### B. Reflexivity of protein-based assignment catalysis offers superior paths to code bootstrapping and optimal gene sequences

The aaRS molecular biological interpreters are the first and, probably the only, products of mRNA blueprints that can implement the translation table embodied in tRNA. Accumulating reflexive genetic information—genes whose expression by rules can, in turn, execute those expression rules—is an intrinsic architectural feature of the PCW that is absent from any RCW. Rapid self-organization of coding in the PCW is driven by reflexive, in-parallel sensing (Fig. 3) of the amino acid phase transfer equilibria that drive folding and thus enable aaRS to recognize both the symbolic information in tRNA (i.e., the syntax) and the chemistry of enzymes (i.e., the semantics) built as interpretations of mRNA sequence information written in the coding language (Fig. 1C).

*The universal genetic code is a nearly unique selection from an inconceivably large number of possible codes and must have been discovered by bootstrapping*. It efficiently maps the chemical properties of amino acids onto the sequence space of triplet codons (Carter and Wolfenden 2016) and is almost ideally robust to mutation. Bi-directional ancestry restricted the tiny fraction of the possible codes that share this optimality (Freeland and Hurst 1998; Koonin and Novozhilov 2009) to an even smaller subset by requiring anti-correlated coding of amino acid physical properties (Zull and Smith 1990; Chandrasekaran, et al. 2013). Discovery of such a rare and highly optimized code through random-sampling natural selection has a vanishingly small probability reminiscent of Levinthal’s paradox about protein folding (Dill and Chan 1997). Far more likely to produce such a result is a series of feedback-accelerated symmetry-breaking processes, like phase transitions, that could bootstrap the earliest prebiotic translation system into existence from some previous, less well-organized chemistry.

The mechanistic implementation (Fig. 3) of reflexivity makes it clear that the requisites for accelerating a bootstrapped discovery of coding are built into the PCW, but absent in the RCW. We envision a minimal, low fidelity instruction set or “boot block” whose realization has been substantially demonstrated ((Martinez, et al. 2015); Fig. 3), and whose feedback-sensitivity could improve itself by elaborating its own resources, much like installing an operating system in a computer at startup. Increasingly specific coding assignments during successive transition steps could take hold only by conferring new selective advantage(s) to the evolving genes, i.e. mRNA sequences, in which they became encoded. This way, such a system could express new meaning in a kind of snowball effect beyond the specific level of fidelity and complexity already achieved.

*The bootstrapping metaphor creates substantially new perspectives on the emergence of genetic coding by integrating local environmental sensing into generating function (Fig. 4).* Coding rules ultimately result from the RNA and/or protein folding rules that generate functional assignment catalysts from sequence. Ribozymal and enzymatic functions are coupled to quite different nano-environmental effects. Specificity, RNA folding depends largely on base pairing because the four nucleotide bases are otherwise almost undifferentiated, having only two sizes and solvent phase transfer equilibria that differ by at most –3.7 kcal/mole in their transfer free energies from chloroform to water (Cullis and Wolfenden 1981). The corresponding phase transfer equilibria of the 20 canonical amino acids (Radzicka and Wolfenden 1988) exhibit approximately five-fold greater variations in polarity and 26-fold greater variation in size. These differences together with the dominance of backbone-backbone hydrogen bonding result in profoundly different protein folding rules.

**Figure 4.**
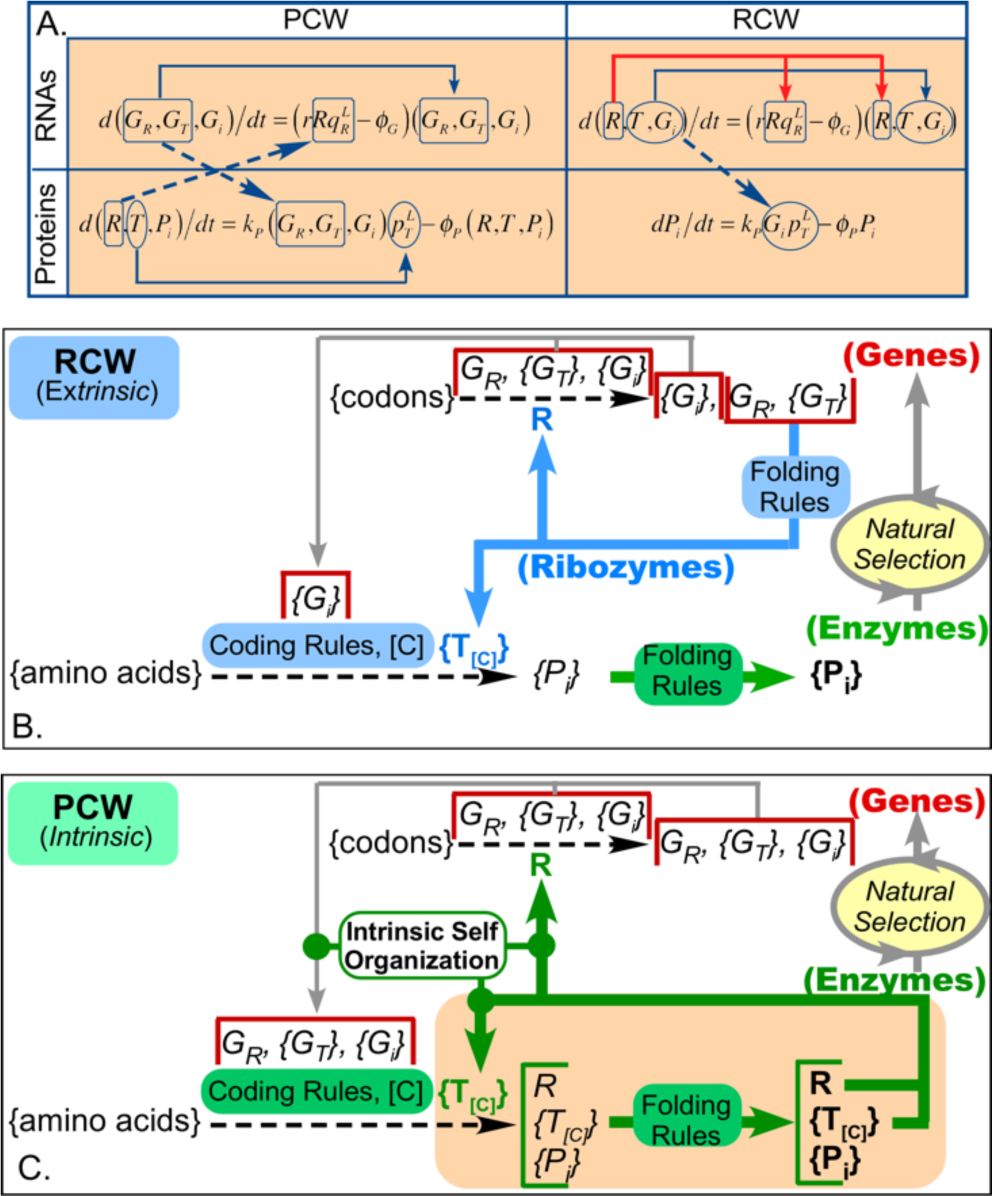
Feedback in riboyzymal (RCW) and protein (PCW) GRT networks. A. Coupled Replicase and Translatase production. Differential equations for gene expression in PCW and RCW are compared for RNAs and Proteins (Wills and Carter 2017). Solid lines indicate autocatalytic acceleration. Dashed arrows form a (hyper) cycle coupling production dynamics of Replicase and translatase in the PCW, but not in any RCW. B. In an RCW, coding rules [C] are implemented by ribozymal assignment catalysts {T_[C]_} that cannot sense the phase transfer equilibria accessible to protein assignment catalysts. Thus, natural selection is the only feedback cycle. Non-aaRS functional proteins {P_i_} furnish the only source of selective advantage, and have no direct influence on the coding rules. C. In the PCW, coding rules are executed by proteins that must first fold. A tighter feedback loop (green arrows) is a structural feature of the reaction network. Protein folding rules determine the function of the assignment catalysts and therefore also the eventual choice of codon assignments, substantially enhancing natural selection.

The differential equations governing expression dynamics (Fig. 4A; (Wills and Carter 2017)) augment the transcendent difference between coding rules derived in an RCW and in the PCW because the synthesis of protein translatases is autocatalytic (horizontal arrows) in the PCW, but not in an RCW. Coding rules in an RCW must be executed by ribozymes (Fig. 4B). An RCW thus cannot provide the intrinsic self-organization necessary to rapidly refine nanospace protein engineering, the essential advantage to be gained from enhanced coding specificity.

Emergence of higher-functioning encoded proteins cannot be triggered by reflexive feedback in an RCW whose expression system itself contains no proteins. Moreover, coding rules based on RNA folding rules are intrinsically insensitive to protein folding rules and/or functionality. Bootstrapping of coded proteins in an RCW would require selecting ever-better *ribozymal* aaRSs through a slow, indirect Darwinian evolution process that could discover protein folding rules only from non-aaRS protein performance. Moreover, ribozymal aaRS variants capable of improved assignments would have to be selected for robustness against mutation in protein-coding genes. The extrinsic self-organization resulting from mutation and higher-level selection in an RCW provides no direct feedback procedure for discovering a translation table that embodies an ordered symbolic encoding of amino acid sidechain chemistry in folded proteins. Such orderliness would have to progressively prove its advantage for the relevant unit of selection, presumably a protocell.

In the PCW (Fig. 4C), coding rules are determined by the catalytic properties of the extant population of protein aaRSs, whose sequences are, themselves, produced by rules acting on the set of genetic blueprints (mRNAs) that encode the aaRS rule executors. This reflexivity in the PCW enables stages of self-organization in genetic coding to occur rapidly, essentially as dynamic phase transitions, because nano-environmental sensing (Fig. 3) can couple the coding rules naturally to protein folding rules. AARS tertiary structures—positioning distant amino acids in primary structure close to one another in space— as determined by amino acid phase transfer equilibria (Fig. 3), furnish the aaRS specificity required to determine the coding rules. Sensitivity of the code to the phase transfer equilibria of amino acid side chains allows those equilibria to feed directly back onto protein aaRS folding and function, naturally producing a refined map of the phase equilibria that govern protein folding and function in the existing code, via the tRNA identity elements (Wolfenden, et al. 1979; Radzicka and Wolfenden 1988; Wolfenden, et al. 2015; Carter and Wolfenden 2016). Thus, in the PCW the mechanism for nanoscale *control* of chemistry, i.e. coding, is determined directly by its *outcome*.

*A PCW also coordinates and optimizes discovery of gene sequences by placing amino acids with different properties in different positions in accordance with their effects on a folded protein.* To consider aminoacylation functionalities as “assignment catalysis” relevant to coding, the specificity for the relevant amino acid must also have become associated with a parallel specificity in choosing primitive “codons” in precursor mRNA. Enhancements that incorporated new amino acids into the programming language had to co-evolve with messages able to exploit them.

This special relationship between amino acid side chain energetics, the coding rules, aaRS sequences, and their genes, establishes reflexivity at an even more fundamental level, acquiring additional bootstrapping from the chemical effects of amino acids occurring in particular relationships to one another in the three dimensional architecture of folded synthetases (Fig. 4C). In other words, a PCW automatically pressures an evolving code to discover and refine the partition between amino acids that gives the genetic representation of functional properties best adapted for survival: an error-minimized code in which amino acids with similar chemical properties are assigned to similar codons. This argument extends to every stage of code expansion. Thus, code evolution in a PCW will inevitably target both near-optimal folded protein functionality and an encoding that represents survival fitness as precisely as possible.

For these reasons *de novo* emergence of genetic coding into a peptide/RNA world appears to have introduced such overwhelming influence on the choice of codons best able to represent the effect of an amino acid entering the developing ecology inside a folding protein that it must be seen as enormously more probable than coding emerging in an RNA World.

#### C. Fidelity: Any simple PCW taking over a more sophisticated ribozymal coding will increase the overall error rate, degrade fitness, and hence be eliminated by purifying selection

The PCW is rooted in phylogenetically-based ancestors capable only of the simplest coding assignments—perhaps one or at most two bits—and consequently also in a coding system necessarily operating at high error rates. Reducing error rates in both replication and translation must certainly have required larger alphabets. To be selected, the functionality of such primordial coding must already have exceeded that of whatever preceded it. Its simplicity appears to rule out scenarios involving proteins “taking over” catalytic functions from any pre-existing world of sophisticated RNA catalysts. For clarity, we henceforth refer to executors of assignment catalysis as RNA or protein “translatases”, to distinguish them from contemporary aaRS.

Any coding system depends on the maintenance of a population of templates that either specify the sequences of ribozymal aaRSs or encode the sequences of protein aaRSs. In an RCW all such templates are required somehow to survive, essentially as parasites, in a world of RNA replicators. <u>A</u> ribozymal coding system, consisting of only ribozymal translatase species, could be functionally autonomous. However, the attractor state of a hybrid ribozymal/protein aaRS system is one in which the protein population also contributes to the overall rate of translation of any genetic template, and more importantly, to its overall error rate.

A separate paper (Wills and Carter 2017) treats this problem in an extension of earlier mathematical models of coding self-organization (Bedian 1982; Wills 1993; Wills 1994; Bedian 2001; Wills 2004) by considering the dynamic stability of co-existing ribozyme- and protein-operated assignment catalysts. We confirm analytically the intuitive conclusion that translation errors would inevitably be higher for any hybrid coding situation driven simultaneously by separate ribozymal and protein translatases than they would be for an optimized system with only one type of aaRS. If both types of translatases effect codon-to-amino acid assignments at different characteristic rates and accuracies the hybrid system will necessarily operate at intermediate error rates. As Eq. (25) of (Wills and Carter 2017) makes abundantly clear, introducing any significant population of protein translatases intrinsically less accurate than an extant ribozymal coding apparatus will undermine the role of the ribozymal translatases, possibly threatening the protein domain with extinction as the selective advantage of ribozymal translatases, indirectly conferred by protein functionality, is diminished. The problem will be extreme in the presence of rudimentary ancestral protein aaRSs that operate a low dimensional translation table. The accuracy of the protein translatases would then be necessarily be very much less than that of the extant ribozymal population, making survival of proteins dependent on the elimination of the protein translatases. Either way, the only path to current molecular biology thus appears to require protein aaRS genes to emerge in concert with other essential encoded protein genes. That requirement highlights the problems arising from coordinating inheritance with gene expression. We therefore turn our attention to the dynamics of template replication and its effect on the evolution of translation.

*Mixed ribozymal and enzymatic protein replicases.* Copying of genetic information lies at the heart of Darwinian evolution. Consider the advent of a protein replicase in a functional RCW in which relatively sophisticated and accurate information copying has evolved through selection of a general ribozymal replicase. Introducing a protein replicase in an RCW generates the same problem just described for the advent of protein translatases. Any protein replicase less accurate than the ribozymal replicase, which is to be expected for the first such proteins to emerge in an RCW, would diminish the probability of correctly copying all genes, including that coding for the ribozymal replicase. Since the system evolution has been optimized under the constraint of the ribozymal replicase’s performance, the system will be at risk of an error catastrophe unless selection purges it of the emergent protein replicase.

*The evident simplicity of the earliest coding apparatus in the PCW poses an insuperable barrier to takeover of any more sophisticated coding apparatus in an RCW*. Newly emerging protein-based assignment catalysts must have been far less specific than the pre-existing ribozymal assignment catalysts envisioned, for example, by Wolf and Koonin (2007), and cannot have been selected within an advanced RCW because their very rudimentary functionality would corrupt any pre-existing ribozymal translation system of higher specificity and diversity. No hybrid set of protein and ribozymal aaRS and/or replicases can have superior fitness to those of a pre-existing RCW, as reinforced by the following considerations:

(i) The more sophisticated the pre-existing RCW, the harder it would have been for early stages of PCW code development to compete. Conversely, the detailed inversion symmetries arising from bi-directionally coded gene (§I) all point to their key role in enforcing differentiation early in the evolution of the genetic code, when it was most vulnerable to parasites with incorrect specificities.

(ii) The dramatic rate acceleration by aaRS protozymes on the other hand, represents a decisive selective advantage in a pepide•RNA world, first by harnessing the chemical free energy transfer of NTP utilization and then by providing a flow of activated amino acids.

(iii) RNA sequences destined to evolve into genes once an accurate translation system had evolved would have had no obvious selective advantage unless the emergent PCW code was practically identical to that operating in the RCW.

Thus, even were an RCW to have existed, it would be irrelevant to contemporary biology if the PCW had to recapitulate the entire genesis of the code. Nor, of course, does any evidence remain of such ribozymal amino acid activating catalysts, or, indeed of ribozymal polymerases. Finally, if the branching phylogenies of protein aaRS provided opportunity for self-organising quasispecies bifurcations, and their evident reflexivity greatly accelerated the search for an optimal code, then, an extensive phase of ribozymal protein synthesis no longer fills any theoretical deficiency in accounting for the genetic code. Thus, it is our view that nature did not re-invent its “operating system” (Bowman, et al. 2015).

#### D. Efficiency: avoiding dissipations leading to Eigen and Orgel error catastrophes

The bootstrapping requirement (§II.B) and the instability of hybrid coding assignment systems with substantially different error rates (§II.C) may reflect inherently complementary arguments for efficient coupling between self-organization of information storage (replication) and readout (translation). Progressive mutational loss of reflexivity leads to progressive increases in the coding error rate (Wills 1994), resulting in the dissipation of free energy flows and ultimately in what have been called “error catastrophes”. Error rates impede self-organization at multiple levels. We examine here the possible coupling between error generation during replication and translation.

*Studies of gene-replicase-translatase (GRT) systems reveal gene replication and coded expression are interdependent.* Are both self-organizational processes so strongly coupled that they emerged simultaneously? Such coupling is not only possible (Smith, et al. 2014) but it occurs spontaneously (Füchslin and McCaskill 2001; Markowitz, et al. 2006; Wills, et al. 2015). GRT systems are intrinsically spatially self-organizing, and unlike the hypothetical RCW no extrinsic, higher level units of selection— i.e. compartmentation—are required to assure their survival.

Living systems now produce proteins from information encoded in genes using aaRS translatases whose genes are copied using protein polymerases. The dynamics of the RNA domains of the PCW and RCW (Fig. 4A; (Wills and Carter 2017)) make it evident that gene and protein production in the PCW are tightly coupled through the population variables that represent the genes and replicase enzyme. Furthermore, translatases in the protein domain are cooperatively autocatalytic.

In contrast, events in the RCW protein domain have no effect on the value of any RNA domain variable, so the dynamics of replication and catalyzed coding assignments are completely autonomous in the RNA domain of the RCW. Moreover, the protein domain is utterly dependent on the RNA domain through the variables that represent the populations of encoding genes and the accuracy of the ribozymal translatase population. It is hard to envisage how any selection pressure that proteins might exert on the sequences of nucleic acids in an RNA World could mold a refined, chemically ordered system of genetic coding.

On the other hand, overcoming the double risk of Eigen- and Orgel-like error catastrophes in information storage and readout implicit in highly coupled molecular biological systems seems equally impossible, until one takes account of the fact that natural selection is a self-organizing force that staves off the potential error catastrophe that threatens information storage (Eigen 1971). Likewise, coding self-organization (Wills 1993) staves off the potential error catastrophe in translation (Orgel 1963). Neither system can be expected to operate unless each limits deleterious effects of the error rate of the other.

Just as power transfer in dissipative electronic structures is optimal if input and output impedances match, so molecular biological organization observed in life’s informational systems may have evolved most efficiently by matching improvements in error rates for nucleic acid replication and protein synthesis at successive development stages. If so, gene expression and replication by functional protein replicases could not have emerged efficiently from a world in which either function was already performed at a higher level by ribozymes.

Direct bootstrapping of genetic information and encoded functional proteins is far more plausible than any scenario in which there was an initial RNA World by three criteria—reflexive feedback, degraded specificity in hybrid systems, and the need to match the complexity of coding to that of protein function. Significant traces in the structural biology of the contemporary aminoacyl-tRNA synthetases therefore suggest that the evolution of genetic coding proceeded by precisely such a sequence of phase transitions, each entailing bifurcations of the information-processing characteristic of the previous stage.

*Impedance matching argues for coevolution of replication and translation.* Fig. 5 illustrates a mechanism coupling biological information storage and readout (dashed lines in Fig. 4A; (Wills and Carter 2017). We conjecture that progressive increases in the dimension of the codon table, *n_eff_*, enhance coding evolution efficiency by matching noise in genetic information maintenance (replication errors and quasi-neutral drift in sequence space) to that from the translation error rate. To paraphrase from a recent definition of “information impedance matching” of information sources to receivers in a different context (Martin 2005), reading out genetic information with as little dissipation as possible requires readout machinery with approximately the level of noise present in the information sources. If errors in either process are either too high or too low, the system will dissipate energy unnecessarily, reducing the readout efficiency. In other words, at any evolutionary stage of developments in molecular biology, the selective effect of the “replicases” and the fidelity of the “translatases” (and any associated accessories) need to limit noise to comparable levels in order to optimize the efficiency of information transfer at that stage.

**Figure 5.**
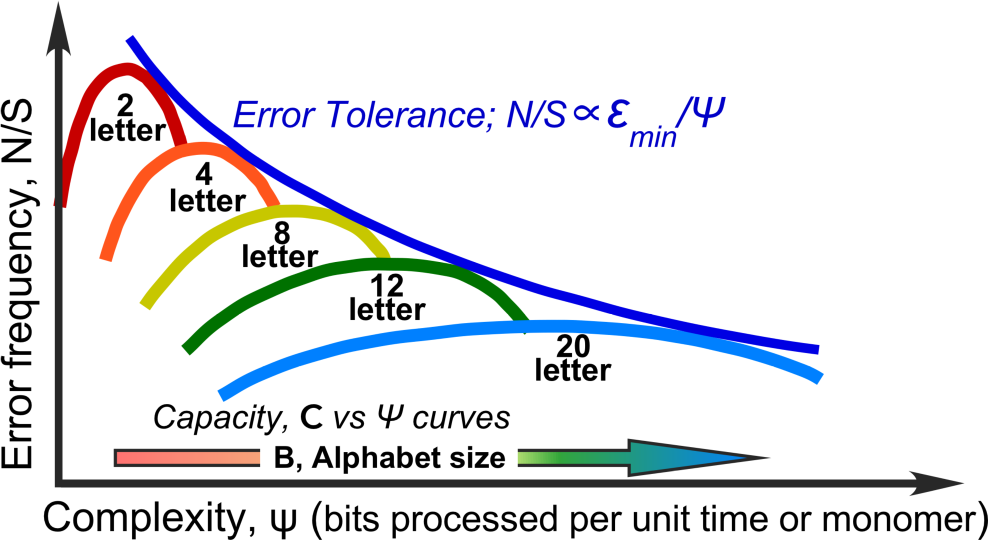
Impedance-matching eases elaboration of coding from a 2-letter amino acid alphabet to a full 20 letter alphabet. Noise, *N*, in the genetic signal, *S*, on the y axis, serves as the primary obstacle opposing information transfer in translation. Increasedcomplexity, on the x axis, must be accompanied by reduced error rates. The error tolerance curve is a hyperbola in which the product of error frequency by complexity, N/S*Ψ, equals the minimum cost of an error, ε_min_, as estimated by Schneider (Schneider 2010). By analogy to the gears on a bicycle’s derailleur, enlarging the alphabet size increases coding capacity, providing a series of matches with the hyperbolic bounding error tolerance curve (dark blue), easing the path to increased fidelity by enabling stepped increases in coding capacity and complexity.

The notion of impedance-matching is well-established physics. Our heuristic reference to it here is supported by the following observations. Error rates appear to be a valid metric for emerging biological complexity over quite large timescales (Lewis, et al. 2016). Specific aspects of that work support this view: (i) Michaelis Menten parameters for the LeuRS and HisRS2 urzymes (Carter, et al. 2014; Carter 2015) suggest that, whereas they are quite impressive catalysts, their specificities for cognate amino acids are well below those necessary to stabilize populations of full-length aaRS, which have much higher fidelities. (ii) Structural studies of the TrpRS urzyme show that its high rate acceleration arises from what appears to be a molten globular ensemble (Sapienza, et al. 2016). In other words, it is a less complex molecule—in a higher entropy state—than a properly folded protein. (iii) The million-fold rate accelerations of both wild type and designed Class I and II protozymes (Martinez, et al. 2015) suggest that the manifold of catalytically competent polypeptides is far larger than previously thought possible. (iv) Presumptive error rates for the aaRS constructs therefore exhibit a monotonic decline with increasing mass, and by implication, increasing complexity.

Moreover, errors quite literally (Gladstone 2016) slow the accumulation of information and hence the growth of complexity in many situations. Thus, although the notion of impedance-matching requires further development in evolutionary theory, including incorporating metrics of “efficiency”, it appears that natural selection and self-organization provide efficient coupling between information storage (replication) and information readout (translation), as if the two processes were impedance matched.

## III. Scenarios for early aaRS speciation and co-evolution of replication and readout

Phylogenetic ancestries of contemporary Class I and II aaRS project convincingly back to a single gene. The simplicity of such a gene furnishes a conceptually consistent “boot block” (Fig. 3) substantially reducing the challenge of understanding how genetic coding might have emerged from a peptide/RNA partnership. Moreover, the detailed inversion symmetries help to explain how such a gene would enforce the initial differentiation necessary to break the powerful forces that make quasispecies centroids strong attractors, substantially strengthening arguments that no genetic code could have preceded the earliest coded protein aaRS. Dual-coding genetic quasispecies exemplified experimentally by the protozyme gene described by Martinez, et. al. (2015) and the urzyme gene proposed by Pham, et al (2007) are thus presumptive ancestors to both Class I and II aaRS superfamilies and the universal genetic code itself.

### A. Why do established protein phylogenies suggest late aaRS speciation?

The strongest argument that an RCW preceded the emergence of proteins is that multiple sequence alignments of contemporary protein families (Aravind, et al. 2002; Leipe, et al. 2002; Koonin and Novozhilov 2009; Koonin 2011), suggest that aaRS diverged late in the succession of protein functions. However, takeover of a ribozyme-based computational translation process must lead in a plausible way to, the observed phylogeny of contemporary aaRS superfamilies.

We believe the conclusion that aaRSs developed after the advent of fully functional proteins based on an alphabet of 20 amino acids rests on two questionable phylogenic assumptions: (i) that domains (~250 amino acids) are the basic unit of remote protein evolutionary history, and (ii) that the evolution of Class I and II aaRS proceeded from independent ancestries. The former assumption fails to account appropriately for the highly mosaic nature of contemporary proteins (Pham, et al. 2010; Li, et al. 2011). The latter ignores the bi-directional coding ancestry of Class I and II aaRS urzymes and protozymes, for which experimental evidence is now exceptionally strong (Pham, et al. 2010; Li, et al. 2011; Li, et al. 2013; Carter 2014; Carter, et al. 2014; Carter 2015; Martinez, et al. 2015; Carter 2016, 2017).

The low fidelity of aaRS urzymes implies that they represent an important, but early stage in the evolution of complexity and hence that deep phylogenies based on aligning intact contemporary aaRS sequences (Aravind, et al. 1998; Wolf, et al. 1999; Leipe, et al. 2002; Wolf and Koonin 2007) are probably misleading, especially in the case of the pre-LUCA heritage of the aaRSs themselves (Wolf, et al. 1999; Wolf and Koonin 2007). Notably, neither domain database (SCOP (Murzin, et al. 1995; Andreeva, et al. 2008); CATH (Pearl, et al. 2003)) has been compiled at sufficiently high resolution to identify the Class I and II urzymes as ancestral forms. Large insertions within the catalytic domains of aaRS were likely accumulated segmentally, from exogenous genetic modules with their own previous ancestry (Pham, et al. 2007), subsequent to their initial evolutionary speciation. Such mosaicity in the multiple sequence alignments, akin to horizontal gene transfer (Leipe, et al. 2004; Soucy, et al. 2015) albeit in shorter segments than those considered by Wolf, et al. (1999), could obscure deeper ancestral evolutionary trajectories involving the urzymes.

Fig. 6 illustrates an alternative phylogeny of Class I aaRS that resolves what we feel to be a mistaken conclusion that the Class I aaRS diverged late in the evolution of the proteome. The scheme traces Class I and II aaRS ancestries from a single gene by two distinct processes—speciation of the bi-directional gene (**I**) and strand specialization to transcend its limitations (**II**). It furnishes a satisfactory account of the increase in structural multiplexing and independent parallel evolution of insertion elements and anticodon-binding domains during a period in which protein synthesis operated with a gradually increasing alphabet size that ultimately required development of editing domains (**III**) to achieve the requisite fidelity of the contemporary proteome.

**Figure 6.**
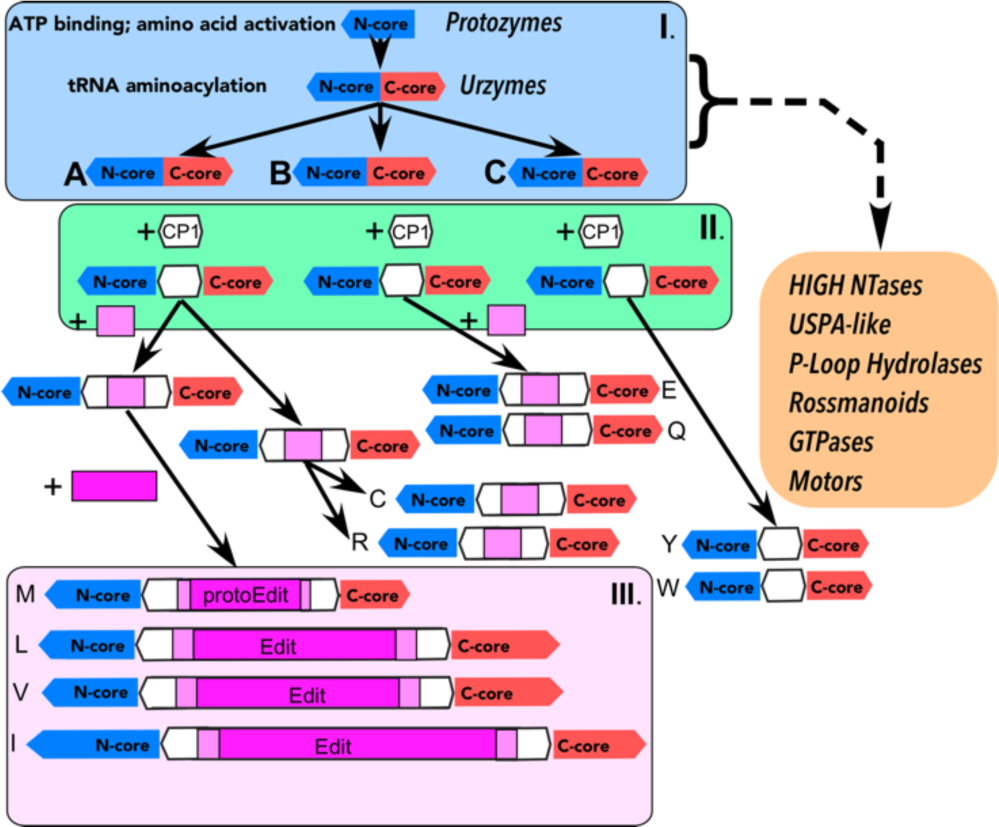
Alternative evolution of Class I aaRS catalytic domains consistent both with phylogenetic analysis and the more ancient ancestry of the most highly conserved modules in the two aaRS Classes. This scenario re-defines the Class I CP1 insertion between the N- and C-terminal modules, N_core_ (blue) and C_core_ (red), of the Class I Urzymes, both of which are portrayed as ancestral to all Rossmannoid superfamilies (Adapted from Fig. 4 of (Aravind, et al. 2002)). The initial CP1 insertion (white) is the origin of most subsequent elaborations of the Class I catalytic domains that appear to have provided the requisite increases in specific amino acid recognition (Carter 2015). Idiosyncratic Class I anticodon-binding domains are and not considered here. We distinguish three phases of aaRS evolution: **I**. Bi-directionally coded with Class II; limited diversity; **II**. CP1 enforces strand specialization; **III**. Hydrolytic editing enhances specificity.

### B. A plausible scenario for co-evolution of information storage and gene expression

We highlight how conclusions from §II change how we think translation might have emerged, and outline a plausible scenario for the co-emergence of information storage and readout. Our scenario is compatible with a rough “impedance matching” in which high noise initially permits co-option of quite unrefined functionalities that comprise groupings of related effects averaged over large but separate regions of sequence space. Noise is gradually brought under control by refining functionalities with distinguishable specificities and selection of genes encoding them, enabling structural diversity and complexity to develop simultaneously with increases in the dimension of the codon table (Fig. 5). Although aspects of this scenario resemble previously outlined marginal scenarios (Martin 2005), its scope, continuity, and its logical, experimental, and phylogenetic support are assembled here for the first time.

The origin of contemporary translation was most likely an intimate co-evolutionary process involving both polymer classes (Carter and Kraut 1974a; Carter 1975). The chief arguments expressed in the following two sections must remain hypotheses until experimental investigation, perhaps guided by ideas we have expressed, can convincingly establish or rule them out. Our recent use of protein design and modular engineering in the experimental colonization of the void that previously existed between pre-biotic organic chemistry (Patel, et al. 2015; Sutherland 2015) and the Last Universal Common Ancestor (Forterre, et al. 2005; Wong 2005; Xue, et al. 2005; Fournier, et al. 2011; Fournier and Alm 2015; Wong, et al. 2016) argues that such experimentation can now be fruitful on a larger scale.

Arguments developed in §II.D imply that replication and translation are necessarily more tightly coupled than is envisioned in the RNA World hypothesis by the need for informational impedance-matching. The overriding challenge associated with the emergence of the genetic code is to develop a scenario in which prebiotic chemistry produces biology through cooperation between nucleic acids and proteins (or their precursors), reflexively, in improving both inheritance and function. Details of such a scenario lie beyond current resources; it is nevertheless appropriate to outline aspects that point toward further work.

*Soon after discovery of catalytic RNA, structural complementarities were identified between extended polypeptide secondary structures and nucleic acids (Carter and Kraut 1974b; Carter 1975; Church, et al. 1977; Warrant and Kim 1978).* The short lengths of polymers required to form such complexes—6-8 amino acids and less than half a turn of RNA double helix, suggested they might have been more stable if their polypeptide and polynucleotide components formed hairpins (Berezovsky, et al. 2000). Their stability as complexes appeared to depend largely on their complementary van der Waals surfaces.

*Stereochemically templated cross-catalysis plausibly accounted for the simultaneous appearance of coding and catalysis*. Helix radii of RNA and double-stranded extended peptides produced optimal van der Waals contacts between the two components at precisely the integral, indefinitely repeating stoichiometry of two amino acids per base (Carter and Kraut 1974b; Carter 1975). This, coincidental, integral stoichiometry enabled a putative rudimentary stereochemical coding. Moreover, specific hydrogen bonding between carboxyl groups of antiparallel β-polypeptide double helices and RNA 2’ OH groups oriented the 3’ OH group as a likely nucleophile suggesting that, in addition, the polar interactions might exhibit templated cross-catalysis, one polymer accelerating the elongation of the other.

*Successive inverted repeats of complementary polypeptide•polynucleotide complexes increased their lengths from ~12 to ~23 to ~46 amino acids and from ~3 to ~6 to ~12 base pairs.* Peptide sequences capable of amino acid activation would have been a plausible consequence, for which we have only circumstantial evidence. Peptides at least 46 amino acids produced by stereochemical coding based on complementary van der Waals surfaces of peptide and RNA backbones might already have begun to exhibit ATP dependent carboxyl group activation, potentiating assembly of peptides. Ligation might then have assembled the first protogenes and a proto-ribosome. Partial complementarity of the 5’ and 3’-terminal halves of the Class I protozyme gene (Carter 2015) suggests coding of the bi-directional protozyme gene by an ancestral RNA hairpin. This reasoning suggests that polypeptide catalytic activities could have preceded even rudimentary genetic coding (Kamtekar, et al. 1993; Moffet, et al. 2003; Patel, et al. 2009).

*The wobble effect (Crick 1966) implies that bi-directional coding would have required a triplet code to provide more than 4 codons*. We encounter here a substantive broken symmetry. The protozyme gene (138 bases) is 6-fold longer than whatever putative RNA hairpin might have been associated with the earliest ~46-residue peptides arising via stereochemical coding. Assuming that such a system could have sustained reproduction nevertheless leaves us with a six-fold gap between the relative stoichiometries of templated cross catalysis and the first true gene expression. Transitions from an initial state in which protein synthesis is initiated without information-bearing genetic templates is envisioned in the theory of coding self-organization and GRT systems (Bedian 1982; Eigen, et al. 1988; Wills 1993). What sort of continuity might have connected an earlier direct, stereochemical coding to indirect, symbolic coding by introducing messenger RNA and the use of adaptors to give the messages meaning?

*The earliest indirect coding emulated the direct stereochemical coding arising from complementary van der Waals surfaces of peptide and RNA backbones.* Analysis of aaRS recognition elements in tRNA acceptor stem and anticodon bases highlighted the capacity of tRNA acceptor stems to encode the size and β-carbon branching, but not the hydrophobicity of amino acids (Carter and Wolfenden 2015, 2016). These properties are necessary and sufficient to encode peptides with the most important characteristics—β- branched side chains favoring extended β-structure and alternating small/large side chains allowing van der Waals access on one face—for assuming structures complementary to the RNA minor groove (Carter and Kraut 1974b; Carter 1975). Symbolic coding by the tRNA acceptor stem could therefore have implemented precisely those features necessary to preserve molecular mechanisms that sustained direct *stereochemical* coding. A selective advantage of that symbolic representation in the proto-tRNA acceptor stem would have been that it smoothed the transition between different stoichiometries—two amino acids per base vs. three bases per amino acid—necessary to implement symbolic coding.

*The ancestral bi-directional gene produced two amino acid activating enzymes, Class I with a modest specificity for larger amino acids, Class II with a similar specificity for smaller amino acids, in keeping with the contemporary specificities of Class I and II aaRS and urzymes.* It is obviously of interest to determine how limited an amino acid alphabet is consistent with catalytic activity of such protozyme genes. Extant experimental results, however, show only that by utilizing the full genetic code the two gene products created from opposite strands can both accelerate amino acid activation ~10^6^-fold. The Class I protozyme possesses a consensus phosphate binding site site (Hol, et al. 1978), suggesting that its catalytic activity may arise from backbone configurations, and not depend entirely on “catalytic residues”.

*The earliest catalysts of aminoacylation may have combined ancestral aaRS with ribozymes (Turk, et al. 2010; Turk, et al. 2011).* It has not been established whether or not the protozymes might also have accelerated tRNA charging with the activated amino acid products. tRNA acceptor stem ID elements likely composed the earliest connection between aminoacylated RNAs and a gene sequence (Schimmel, et al. 1993; Rodin, et al. 1996; Henderson and Schimmel 1997; Rodin and Rodin 2008; Rodin, et al. 2009; Rodin, et al. 2011). Dependence of aaRS tRNA affinity on acquiring an additional, anticodon-binding domain suggests that, in contrast to amino acid activation, aminoacylation may have originated in polypeptide•RNA collaboration and was later taken over by urzymes with the rudimentary capability to recognize tRNA acceptor stems (Li, et al. 2013).

## CONCLUSIONS

The continuing search for ever better RNA replicases (Wochner, et al. 2011; Attwater, et al. 2013; Sczepanski and Joyce 2014; Taylor, et al. 2015; Horning and Joyce 2016) has been no mean achievement. However, we argue here for a more holistic and ambitious set of goals than those fueling that search, anticipating that data and theory in §I, §II, and (Wills and Carter 2017) will stimulate discussion and further research on questions relevant to the origins of gene expression, biology’s readout mechanism. A high degree of coherence connects theories of self-organization to the experimental, structural, and phylogenetic aspects of the evolution of the aaRS enzymes that implement gene expression today.

1. Pronounced inversion symmetries in the amino acid substrates, catalytic residues, tertiary, and secondary structures are evident in phylogenetic, structural, and biochemical data for contemporary Class I and II aaRS and arise from their bi-directionally coding ancestry.
2. Inversion symmetries assure maximal structural and functional differentiation between the two classes, a necessary precondition for their survival in competition with parasitic molecular forms.
3. tRNA identity elements that implement coding efficiently capture the amino acid phase equilibria that drive protein folding and are optimal for bi-directional coding (Zull and Smith 1990).
4. Bi-directional coding and the uniqueness of the coding table create a reflexive feed-back cycle to guide rapid evolutionary emergence of protein aaRS genes by bootstrapping rapidly to an optimal coding table and mRNA sequences. Ribozymal assignment catalysts lack such reflexivity.
5. Hybrid system expression dynamics show that any emerging PCW with a lower-dimensional coding table than that of a pre-existing RCW would necessarily have been eliminated by purifying selection before it had sufficient time to expand the dimension of its coding table.
6. Coupling of dynamic equations for gene-translatase-replicase (GRT) systems suggest that matching of error rates maximized the probability of launching replication and translation.
7. (1)-(6) imply that replication and readout emerged simultaneously from a peptide•RNA partnership. We outline a more probable scenario than an RNA world for the origin of biology.
8. Molecular constructs (§I) enhance the ability to test specific elements of proposed scenarios.

## Acknowledgments

This work was supported by The National Institute of General Medical Sciences, (grant numbers R01-78227 & R01-90406 to C.W.C., Jr.). This publication was made possible also through the support of a grant from the John Templeton Foundation. The opinions expressed in this publication are those of the author(s) and do not necessarily reflect the views of the John Templeton Foundation. H. Fried (Cursor Scientific Editing and Writing, LLC) made many useful suggestions on an earlier draft.

